# Lenticular nucleus volume predicts performance in real-time strategy game - cross-sectional and training approach using voxel-based morphometry

**DOI:** 10.1101/2020.07.20.205864

**Authors:** N. Kowalczyk, M. Skorko, P. Dobrowolski, B. Kossowski, M. Myśliwiec, N. Hryniewicz, M Gaca, A. Marchewka, M. Kossut, A. Brzezicka

## Abstract

It is unclear why some people learn faster than others. We performed two independent studies in which we investigated the neural basis of real-time strategy (RTS) gaming and neural predictors of RTS games skill-acquisition. In the first (cross-sectional) study we found that experts in the RTS game StarCraft II (SC2) had a larger lenticular nucleus volume than non-RTS players. We followed a cross validation procedure where we used the volume of regions identified in the first study to predict the quality of learning a new, complex skill (SC2) in a sample of individuals who were naïve to RTS games (second training study). Our findings provide new insights into how the volume of lenticular nucleus, which is associated with motor as well as cognitive functions, can be utilized to predict successful skill-learning, and be applied to a much broader context than just video games, e.g. contributing to optimizing cognitive training interventions.

## Introduction

Some people learn faster than others. Skill learning - the process that makes people more accurate, efficient and faster in a given task - depends on several personal characteristics. From the psychological perspective, there have been theories and data regarding the prediction of learning based on individual differences in non-cognitive and cognitive determinants since the 1950s. Specifically, skill learning was well-predicted by age (*1*), general ability measures, such as verbal, spatial and numerical reasoning, as well as working memory capacity (*2*-*4*) and fluid intelligence (*5*). Additionally, Yesavage et al. (*6*) showed that individuals with higher mental status in terms of their scores in the Mini-Mental State Examination (MMSE) (*7*) were characterized by better outcomes after memory training. There were also attempts to verify more basic cognitive abilities like perceptual-speed and psychomotor characteristics as predictors of skill development in complex paradigms (i.e. air traffic control simulation task) (*8*). It needs to be pointed out that because the process of skill acquisition is dynamic, cognitive and non-cognitive constructs may differentially determine individual differences in task performance, depending on e.g. the type and stage of the task’s practice (*9*). For example, one of the most well documented associations between individual differences in the non-cognitive domain and skill acquisition were from the personality domain (e.g. Big Five inventory) (*10*).

The picture is even more complicated when it comes to the relationship between the ability to master new skills and its neuroanatomical predictors. Plenty of cross-sectional imaging studies have demonstrated structural brain differences between experts in music, sport, video games and non-experts and showed that experts had more gray matter volume (GMV) in certain brain regions (*11*-*17*). However, deliberate practice is necessary but not sufficient to account for individual differences in experts and novices (*18*-*20*). One of the criticisms of cross-sectional studies as providing the evidence for practice-dependent brain changes is that preexisting differences in brain organization could explain some of the differences we observe between experts and non-experts. For example, the GMV in the hippocampus of London taxi drivers may be larger because they have regular experience with navigation, or because they have some brain structure characteristic that predisposed them to become taxi drivers (*21*). Another study showed complementary evidence in the domain of specific predispositions and experience-dependent brain plasticity (*22*). There are separate groups of studies that assessed regional brain morphometry characteristics of subjects who underwent longitudinal assessments using magnetic resonance imaging (MRI) (*23*). They showed changes in gray matter during skill acquisition of e.g. juggling (*24*), playing video games (*25, 26*), learning languages (*27*), playing music (*28, 29*), aerobics (*30, 31*) and established a theory of brain volume expansion in task-relevant areas as an indicator for neural plasticity (*32*), especially during initial stage of learning (*33*). Given the above described examples, the debate about the predictive neural markers of learning has thus far been inconclusive and still is one of the most challenging areas in cognitive neuroscience. Currently we can observe a growing interest in how individual differences in the structure of the human brain can influence the ability to learn and master complex skills (*34, 35*). Particularly, since the brain’s gray matter characteristics are one of the most adequate biological structures that could determine cognitive abilities, it is essential to look at it as a variable predicting training efficacy.

Scientists reported that pre-existing neuroanatomical profiles, including both cortical thickness and white matter microstructure, predict the outcomes of individuals following multi-strategic memory training (*36*). There is also evidence suggesting that variance in white matter structure correlates with the ability to learn musical skills in non-musicians, offering an alternative explanation for the structural differences observed between musicians and non-musicians (*37*). Only a few studies have explored pre-existing neural characteristics in the case of learning how to play video games, as an example of complex skill acquisition. Momi in 2018 (*38*) identified that lingual gyrus is involved in the ability to predict the trajectories of moving objects in action video games. Other researchers have mostly investigated the volumetric characteristics of the basal ganglia, a group of subcortical nuclei involved in motor and procedural learning, as well as in reward learning and memory (*39*-*41*).

In the study reported here, we examined brain GMV - related differences in the acquisition of skill in a novel and complex cognitive-motor task - the RTS game. We chose SC2 game based on evidence suggesting that playing this cognitively demanding strategic video game requires a host of specialized skills, including translating mental plans into motor movements, performing actions with precise timing, bimanual hand coordination, and processing rapid visual information (*42*). These skills are trained and become more and more automatic with quantity of practice (*43*). What is more, we can use telemetry data from the game (e.g. Perception Action Cycle (PAC) latency, Actions Per Minute (AMP) or Hotkey Selects (HS) usage) to have more detailed measures of skill learning during the course of the game and use it for further investigations. Moreover, a player’s current skill level can be determined by the their position in one of the six tiers in the game (a detailed description is provided in the method section) and, what is also important, it classifies players based on Elo score like rating systems what allows for the objective assessment of changes in player’s expertise over time. One additional benefit of SC2 is that it belongs to a group of games that comprise professional electronic sports (eSports). Because competitive and professional players of eSports titles dedicate a great deal of time to playing individual games, they are a sample with a more stable source of video game experience. This makes the analysis of connections between in-game actions and brain structures more feasible.

In the current study we wanted to test whether it is possible to predict the level of skills acquired during RTS game training based on specific brain GMVs. Our main hypothesis tested the possibility of predicting the quality of skill acquisition (SC2 game) based on the volume of brain regions identified in a group of expert players. In our attempt to understand the neural predictors of learning success in the SC2 environment, we took a two-step approach. First, we analyzed a cross-sectional sample of expert RTS players (those placed in the top five SC2 leagues) and non-players (NVGPs) to investigate whether RTS video game experience is associated with volumetric differences in gray matter. In the second step, we used information gathered during the first study to inform the analyses on data gathered during the second, training study, where naïve volunteers were trained with SC2.

With those steps, we followed a cross validation procedure (*44*) in which one sample of subjects is used to identify brain regions (ROIs) that differ two groups (in our case: RTS experts and NVGPs), and another sample (NVGPs) to predict skill acquisition (in our case: complex skill learning during SC2 training) from the ROIs identified in the first step. Details of this procedure are depicted in Fig. 1A and 1B. To our knowledge, this is the first study where neuroanatomical individual differences between healthy adults were used as a predictor of learning outcomes in an RTS action video game.

**Fig. 1.**
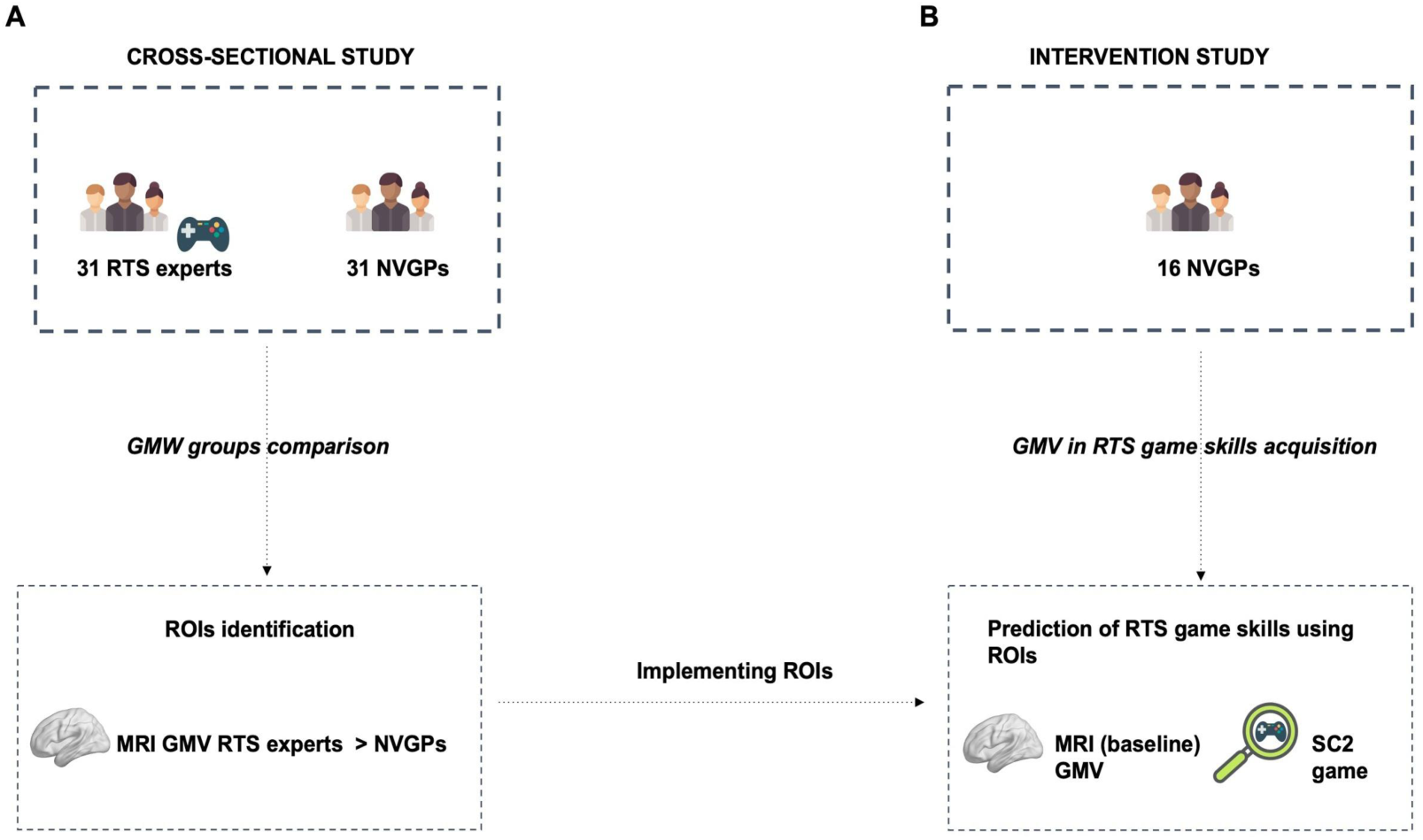
Overview of the study design. First, the GMV ROIs were identified in the cross-sectional study **A**, and then they were used to predict RTS game skill acquisition in independent training study **B**. As a next step a longitudinal study was conducted with the same RTS game as in the first study (SC2) on a new group of non-video game players. Abbreviations: GMV - gray matter volume, MRI - magnetic resonance imaging, NVGPs - non-video game players, ROIs - regions of interest, SC2 - StarCraft II

## Results - cross-sectional study

### Higher GMV in RTS experts compared to NVGPs

Thirty-one RTS experts (in SC2 game) were compared to thirty-one NVGPs using high resolution T1-weighted images (Tw1). Differences between players and non-players in GMV were calculated using whole brain voxel-based morphometry (VBM) analyses. RTS experts had significantly higher regional GMV in the right lenticular nucleus (putamen and pallidum) compared with non-experts (peak MNI coordinates x = 22; y = −11; z = 7; *t* = 5.54; cluster size = 2125 voxels), *p* = 0.04 corrected for multiple comparisons with family Wise Error (FWE) correction at cluster-level using cluster size. The obtained result is in the lenticular nucleus, a structure consisting of the putamen and the pallidum (also commonly called globus pallidus), which are separated by white matter tracts called lateral medullary lamina. In the whole brain analysis, the right lenticular nucleus was the only area showing a significant difference in RTS experts in comparison to NVGPs (Fig. 2 A). There were no significant differences in GMV for the reverse contrast (NVGPs vs. RTS experts).

**Fig. 2.**
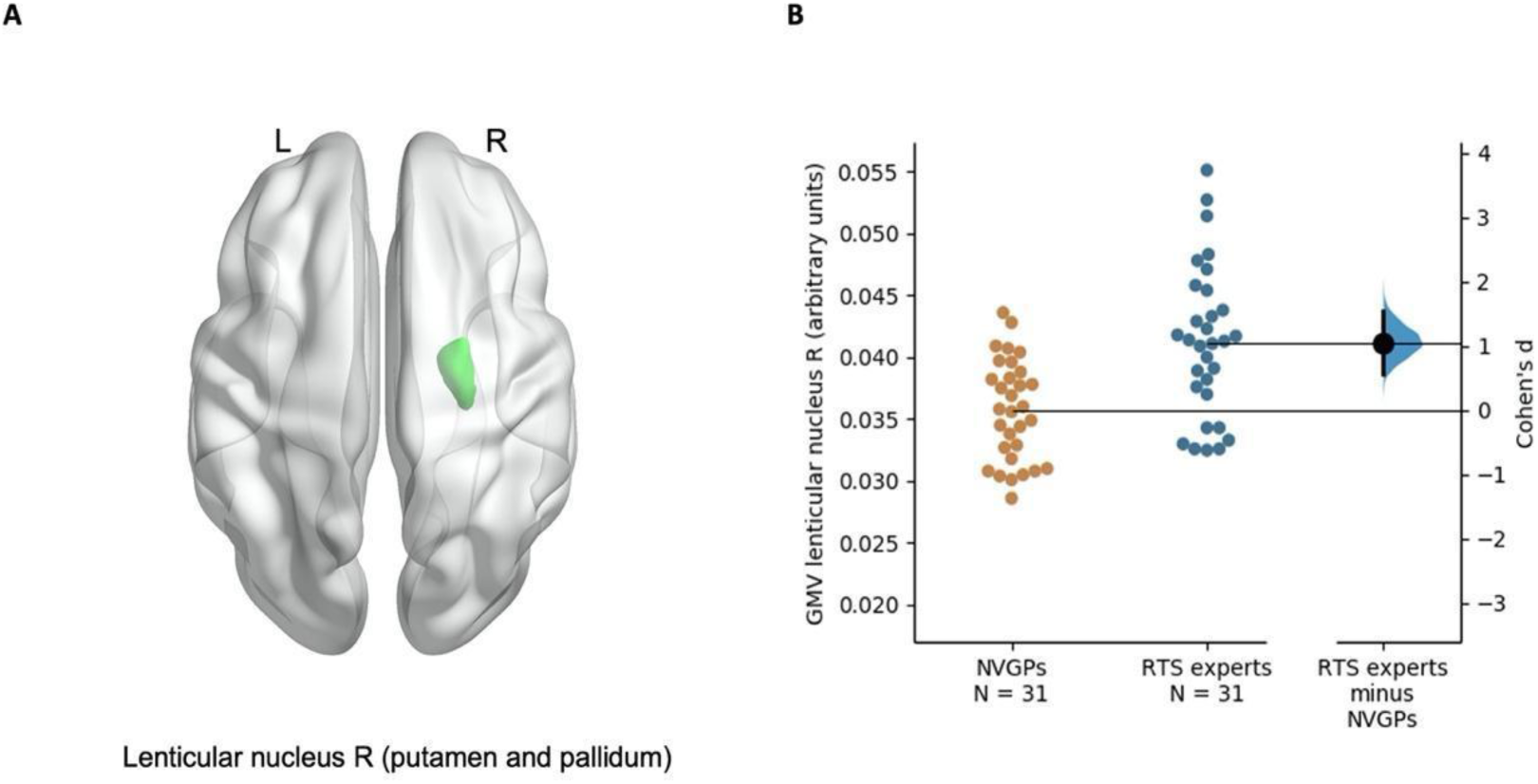
Differences in GMVs between RTS expert players and non-players. **A** Results from VBM y analyses showing RTS experts > NVGPs difference in GMV - part of the lenticular nucleus (peak MNI coordinate x = 22; y = −11; z = 7; *t* = 5.54; cluster size = 2125 voxels), Clusters from the whole brain exploratory analysis using FWE cluster correction. The lenticular nucleus is a collective name given to the putamen and pallidum (also commonly called the globus pallidus); both are nuclei in the basal ganglia. **B** Presentation of differences in GM between RTS experts and NVGPs. Cohen’s d presented to show the effect size of the difference (*d* = 1.06). Abbreviations: GMV - gray matter volume, NVGPs - non-video game players, ROIs - regions of interest, L - left, R - right, SC2 - StarCraft II

We calculated Cohen’s *d* together with power (Fig. 2 B). Cohen’s *d* was 1.061 and using G*Power (*45*) we had 82 percent power to detect differences between groups.

No significant correlation (Spearman’s) was observed between the GMV within the lenticular nucleus and the index of experience in SC2 (hours spent playing SC2, RTS expert group only), *r* = 0.19, *p* = 0.32.

## Results - training study

### Regional GMV as a predictor of RTS game skill acquisition

In the next study we used RTS game - SC2 as a tool to study complex skill learning in a longitudinal setup. We computed the variable indexing the weighted time spent on every level of SC2 difficulty, which reflects performance in the game. Figure 3 presents the time (hours) spent on each level for all participants.

**Fig. 3.**
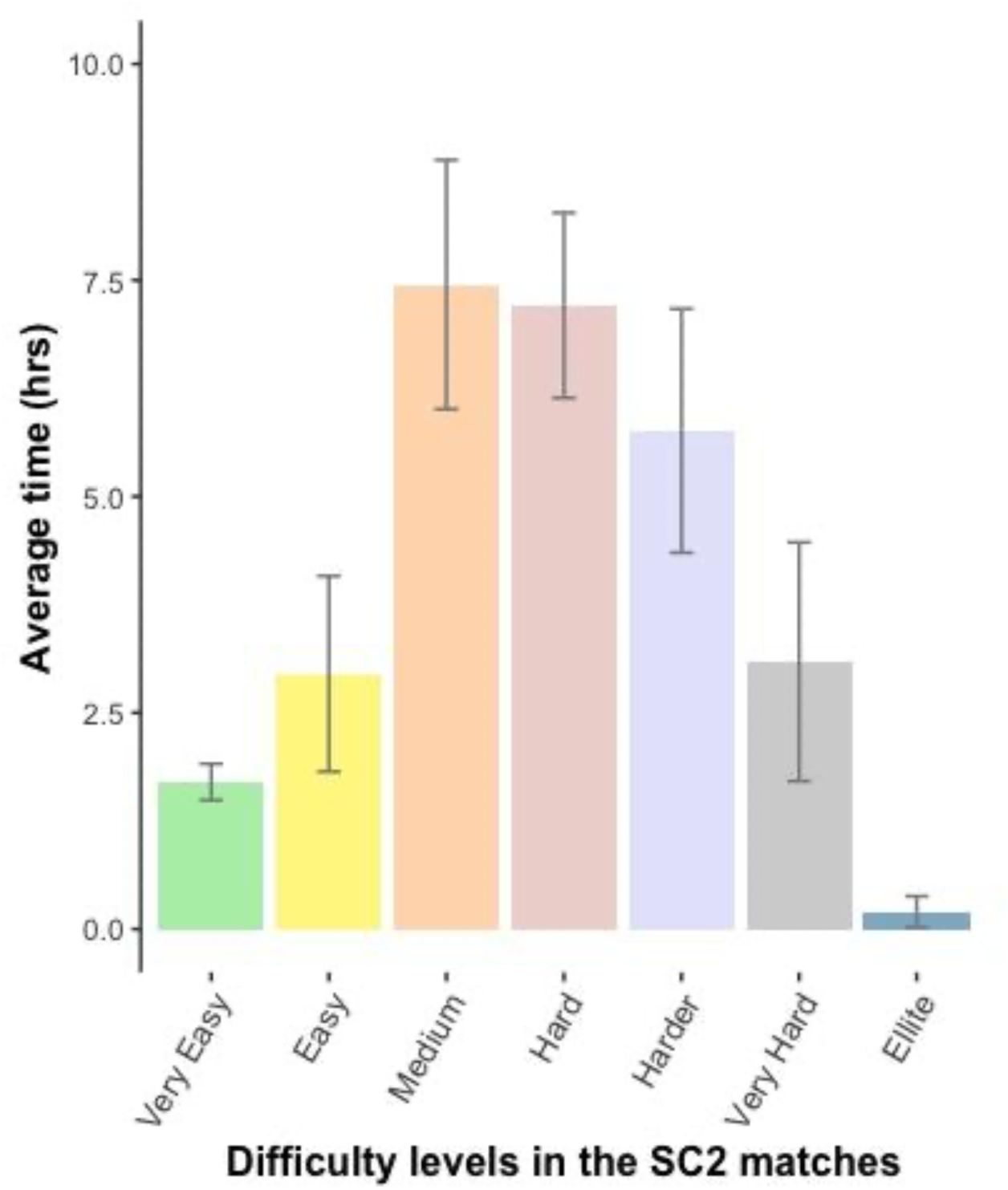
Distribution of the average time (hours) spent on each difficulty level in SC2 for all participants. Presentation of possible difficulty levels in the SC2 matches: Very Easy, Easy, Medium, Hard, Harder, Very Hard, Elite. None of the participants reached the Cheater level, so we included only seven levels. The weighted time spent on each level of SC2 difficulty (the time spent on the second level was multiplied by two, the time spent on the third level by three, and so on) was computed for each participant. The final result is a standardized (group-wise) sum of the time spent on all difficulty levels, which reflects performance in the game. This indicator was used in the correlational analyses. Presentation of average time (M) and standard errors (SE) for number of hours played for each difficulty level in SC2 game. Abbreviations: SC2 - StarCraft II

To specifically target our hypothesis, we employed the ROI analysis method to longitudinal data. Our ROIs were defined based on the results from our cross-sectional study, which showed that SC2 performance was associated with volumes of the ventral striatum (putamen and pallidum).

We used anatomical ROIs based on GMV differences in areas that were related to RTS gaming activity in our first, cross-sectional independent study. We found that the volume of both putamens (left: *r* = 0.67, *p* = 0.01 and right: *r* = 0.57, *p* = 0.02) (Fig. 4 A) as well as both pallidums (left: *r* = 0.62, *p* = 0.01 and right: *r* = 0.62, *p* = 0.01) correlated positively with RTS game skill acquisition (Fig. 4 B) (Spearman’s correlation). No correlation metrics survived the false discovery rate (FDR) p<0.05 correction for multiple comparisons so the results presented here are uncorrected for multiple comparisons. There were no training related changes in the GVM of the examined brain structures.

**Fig. 4.**
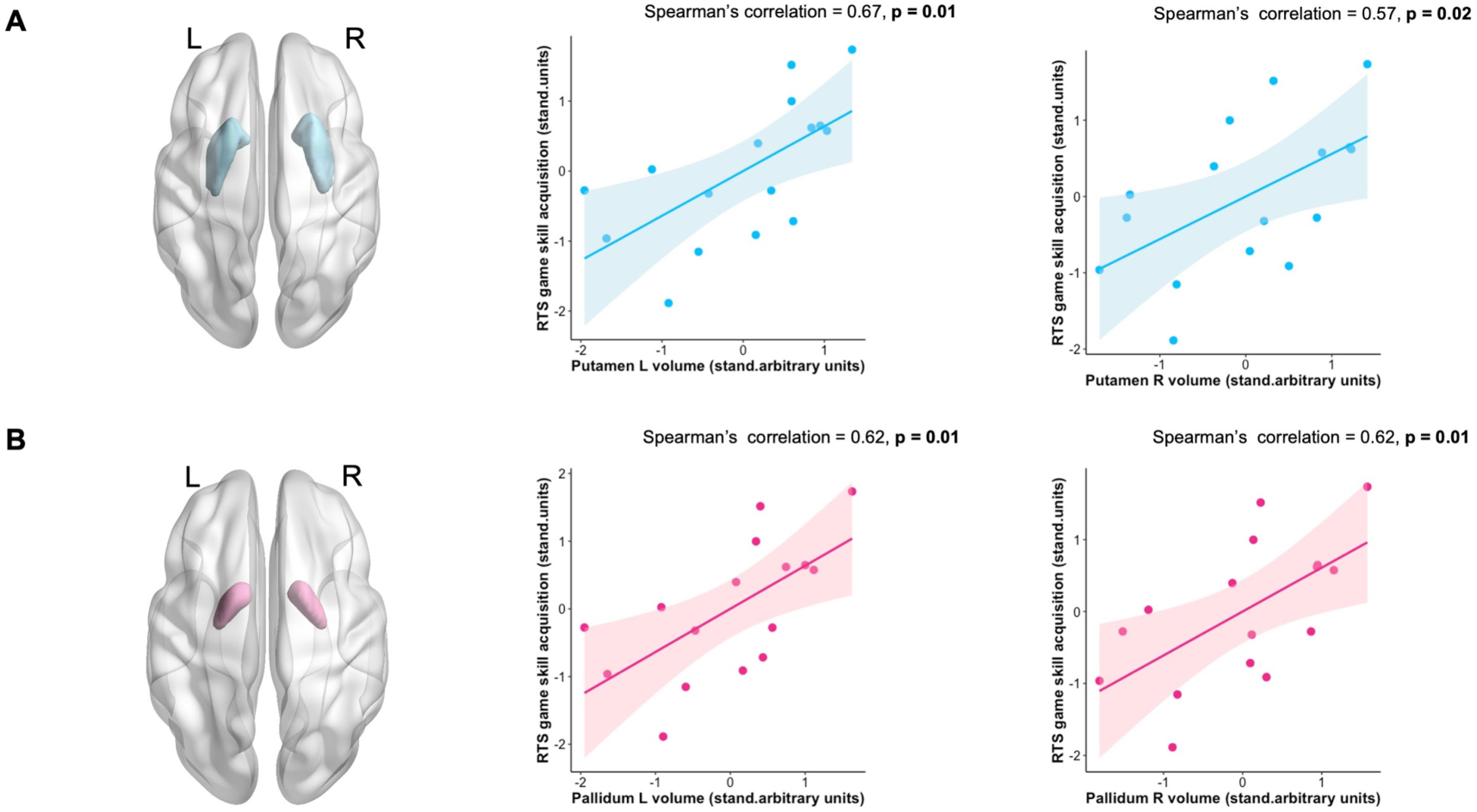
Predefined ROIs (right, left putamen A and pallidum B) and scatter-plots portraying the relationship between GMV in ROIs and RTS game skill acquisition. Based on the significant difference in both putamen and pallidum (part of the lenticular nucleus from Figure 3) in cross-sectional study, we chose those areas as a ROIs for training study to evaluate the patterns of RTS game skill acquisition. Panels show areas with a significant (bolded) positive correlation between mean GMVs in the ROIs with SC2 game performance. The blue color represents the results for the putamen, and the pink color represents the results for the pallidum. The results of correlation analyses are uncorrected for multiple comparisons. Abbreviations: ROIs - regions of interest, L - left, R - right

### Regional GMV and Perception Action Cycle latency in RTS skill acquisition

Our next step was to check what type of in-game behavior correlates with VBM assessments of GMV. Using measures of cognitive-motor abilities extracted from SC2 game replay data from sixteen participants, we constructed three indicators based on in-game actions performed by trainees: [1] PAC latency - time (in milliseconds) from a point-of-view change (switch in focus of attention) to the occurrence of the first action issued by the player (indexing motor reaction). [2] HS usage - expressed as the average number of hotkey presses per minute in each game, where each such action represents an automated selection of multiple units or buildings. Hotkeys are used to aid in the management of dispersed elements of the game. [3] APM - the average number of actions performed during each minute of the game (measure of cognitive-motor speed). We defined PAC latency as a cognitive marker of SC2 expertise, APM and HS usage as a motor markers of SC2 expertise (*42*). We divided the whole training time of each trainee into quartiles and computed PAC latency, HS usage and APM for each quartile (within subject) (*42*).

We found a significant, negative correlation between PAC latency in the first quartile and GMV in all of our predefined ROIs (left and right putamen [left: *r* = −0.58, *p* = 0.02 and right: *r* = −0.43, *p* = 0.10 - tendency level; Fig. 5 A), as well as both pallidums (left: *r* = −0.57, *p* = 0.02 and right: *r* = −0.54, *p* = 0.03; Fig. 5 B). No correlation analysis survived the FDR p<0.05 correction for multiple comparisons so the results presented here are uncorrected for multiple comparisons. Correlations between PAC latency and ROIs volume for the second, third and fourth quartiles were not found. We also conducted correlational analyses for all quartiles for HS usage, APM, and all predefined ROIs, but there were no significant correlations. All correlation coefficients and significance levels are provided in Table 1. PAC latency distribution for each participant is presented in Fig. 6.

**Fig. 5.**
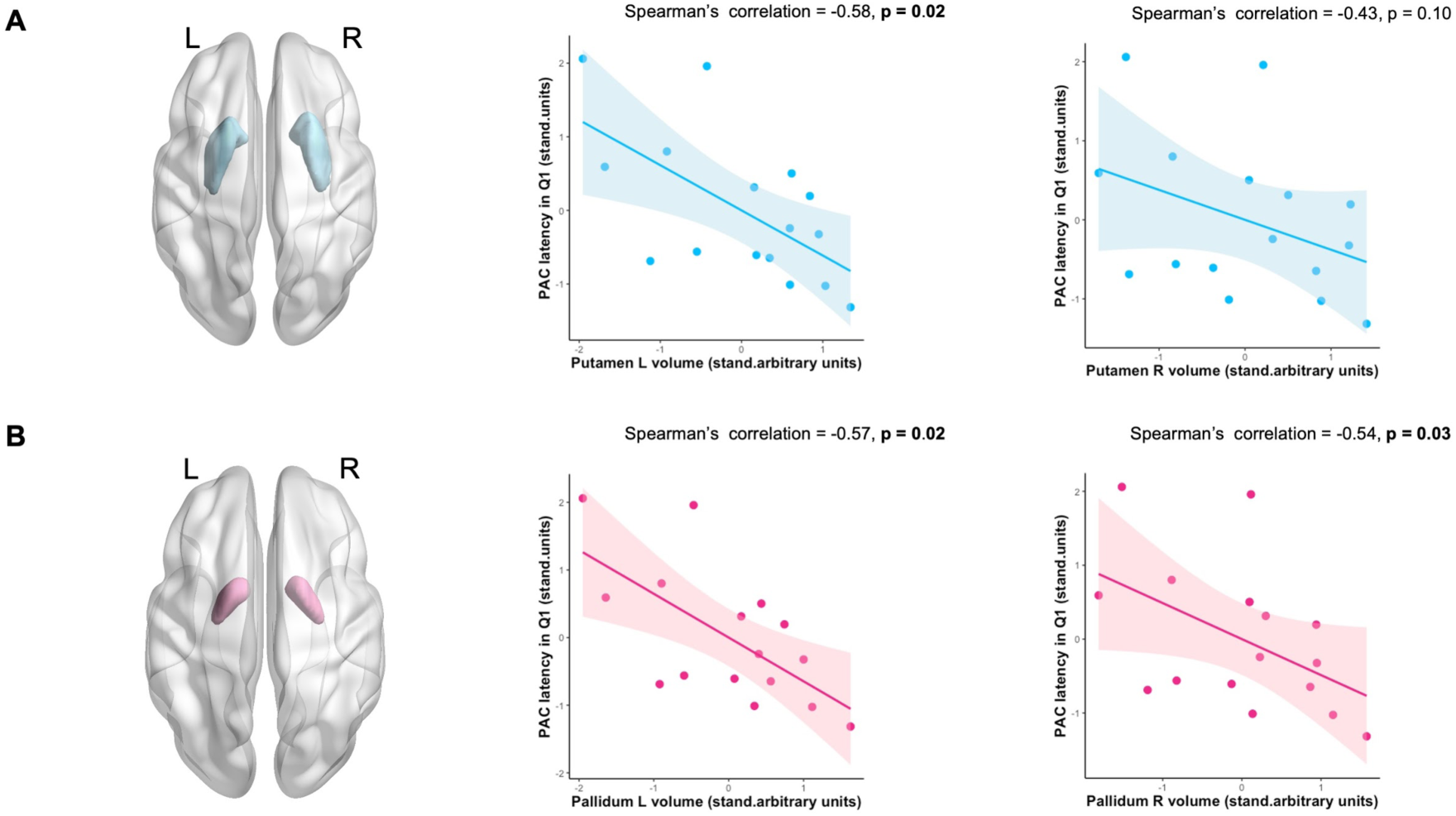
Predefined ROIs (right, left putamen A and pallidum B) and scatter-plots portraying the relationship between GMVs in ROIs and Perception Action Cycle latency in the first quartile. The brain area with a significant difference (part of the lenticular nucleus from Figure 3.) was the datum point to choose the GMVs in the ROIs (both putamen and pallidum) which were included to evaluate the patterns of Perception Action Cycle latency in the first quartile (Q1). The panels show areas with a significant (bolded) negative correlation between the mean GMVs in the ROIs with PAC latency in Q1. The blue color represents the results for the putamen, and the pink color represents the results for the pallidum. The results for correlation analyses are uncorrected for multiple comparisons. Abbreviations: ROIs - regions of interest, PAC - Perception Action Cycle, Q - quartile

**Fig. 6.**
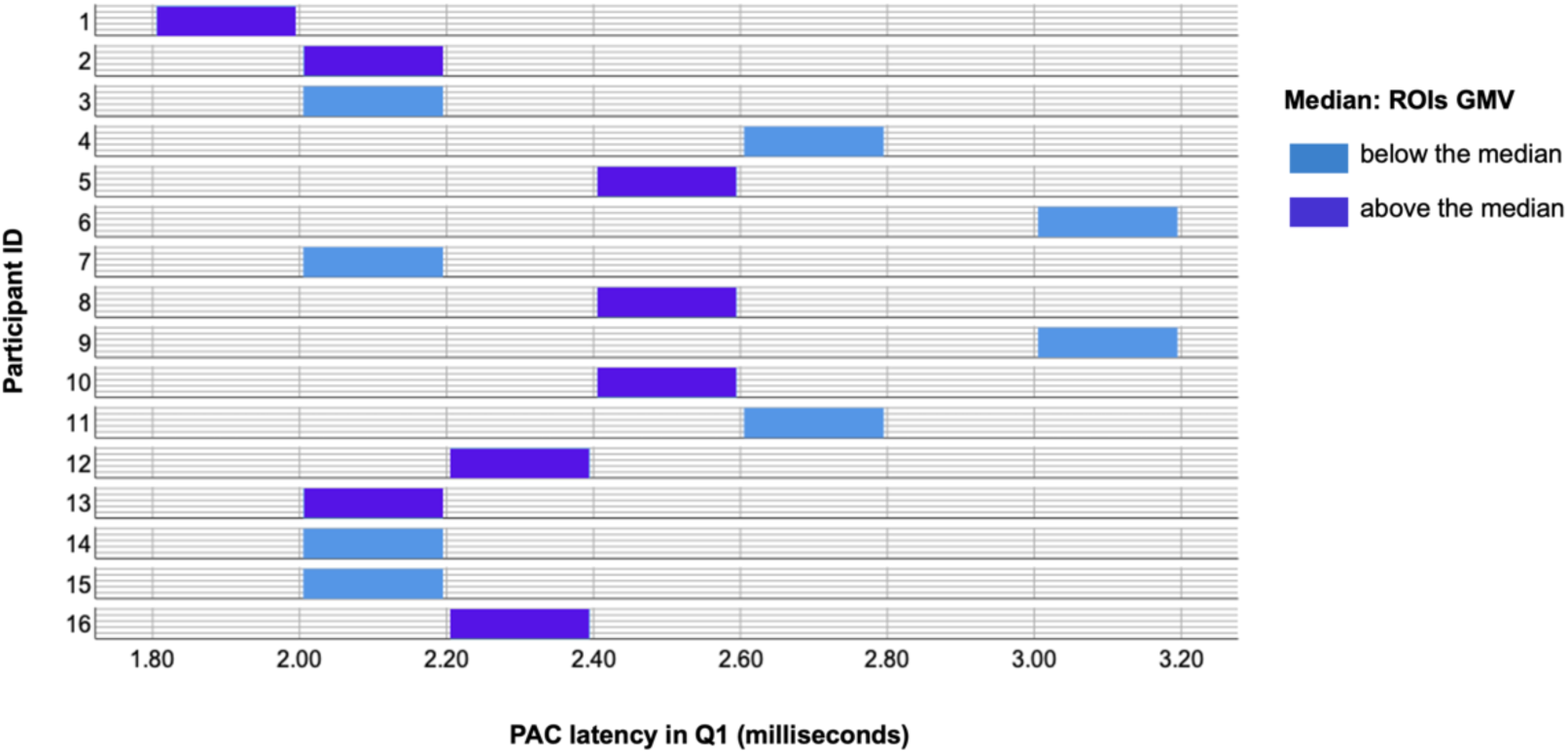
Perception Action Cycle latency distribution for each participant. Additionally, the plot represents the PAC latency distribution in the first quartile (Q1) separately for participants below and above the median: computed from the average GMV for all the ROIs (both right/left putamen and pallidum). Abbreviations: ROIs - regions of interest, PAC - Perception Action Cycle, Q - quartile

## Discussion

In training-related plasticity studies, interindividual differences in learning performance have not received much attention. The large number of publications focused on behavioral improvement and experience-dependent structural changes in the brain. However, neural factors of predisposing to complex skill learning, such as video games acquisition, appear to play an important role in optimizing the training paradigms dedicated to increase the subject’s efficiency and brain plasticity.

In the two studies described here, using VBM, we observed that the volume of the right lenticular nucleus (part of the basal ganglia) was predictive of success in the complex RTS game SC2. Experts in SC2 (players from the top five leagues) had larger basal ganglia compared to people who do not play RTS games. When we took into consideration the volume of the areas identified in the first study in a completely new, unrelated sample of individuals who were naïve to RTS games, we were able to predict the quality of learning in SC2. The regional differences in the volume of the basal ganglia correlated with the pace at which participants learned to play this complex video game. In our opinion, the results presented here provide new insights into how brain volume measurements can be utilized to predict the success of the skill-learning process, and can be applied to a much broader, than just video games, context.

The first, cross-sectional, study showed more GMV in the right lenticular nucleus (putamen and pallidum) in RTS experts when compared to NVGPs. In the second, training study we explored whether the pre-existing volume of the putamen and pallidum can predict improvement in the RTS skill acquisition in novice players. We, in fact, confirmed that the GMV of predefined ROIs was correlated with complex skill acquisition, measured as time spent on more demanding game levels, which is treated here as a proxy of a complex skill learning. The correlation was found in both the right and left lenticular nucleus (putamen and pallidum), whereas in the first study we found differences between experts and non-players in the right lenticular nucleus only. The lenticular nucleus is a subcortical structure within the basal ganglia, comprised of the putamen and pallidum and constitutes a relay station, conveying information between different subcortical areas and the cerebral cortex (mainly primary motor cortex and the supplementary motor area) (*46*). Both pallidum and putamen play an important role variety of motor acts, including sequential motor learning (*47*) and movements control (*48*-*50*) including the operation of a joystick (*51*). It is also abundantly clear that the pallidum and putamen neurons are involved in more than just the organization and/or execution of movements. They are also actively involved in a variety of cognitive functions such as visual attention (*52*), working memory (*53, 54*) and cognitive control (*55*). Additionally, pallidum neurons encode actions such as actual location of the target on a screen, as well as monitor behavioral goals (spatial or object), indicating that this region is involved in goal-directed decisions and action selection (*56*).

To properly understand the results from the two studies presented here, we need to take into consideration the dynamics of the learning process and expertise levels. Playing a demanding RTS game like SC2 requires the engagement of a wide range of cognitive and motor functions (*42, 57, 58*). However, the degree to which each of these functions is engaged is not likely to be equally engaged across all stages of SC2’s learning. Specifically, early attempts to acquire a novel skill, especially as complex as learning to play SC2, are characterized by effortful, explicit information processing which proceeds under executive control functions (especially working memory). As the practice advances, the skill becomes less effortful and more proceduralized, with almost complete automaticity attained at the expert levels of performance (*59*). Our observations paint a pattern of results suggesting that such a process is taking place within the basal ganglia structures. The result of more GMV in the right lenticular nucleus of expert RTS players may stem from effective use of learned motor sequences (especially automatic movements). On the road to success in most RTS games - including SC2 - expertise level is commonly multi-staged, and connected with acquiring a higher and higher degree of automatization of specific sequences of movements. Such automatization allows expert players (like those in our first study) to perform actions that were at first (in novice players, like in our second study) complicated and cognitively demanding, with minimal effort (*60*). SC2 has an economic component, which means that players have to spend resources on the production of military units and structures, and hence many of the player’s decisions / strategies are related to the balancing of expenses on military and economic strength. Secondly, the game board, called the map, is much larger than what the player can see at one time. Thirdly, players do not have to wait for the opponent to play their turn, so the pace of the game is incomparably faster than in e.g. chess. Players who can more effectively and quickly implement their strategy have a huge advantage. Therefore, motor skills, mainly related to handling the keyboard, are an integral part of the game that leads to victory. And thus, the growth of the subcortical structure is probably the result of the above-described experience. We know from other studies that the putamen plays a special role in game related processes, and is also important for movement preparation, learning, and motor sequence control (*50, 61*-*63*). An additional confirmation that video games strongly stimulate motor skills, especially those that are highly specialized and automated, was conducted by Borecki et al. (*64*). Their study assessed a wide range of hand-movement coordination skills and demonstrated that FPS players were able to use motor skills more effectively than control subjects, and the scope of these skills included: improved targeting accuracy, reduced tremors, more effective eye-hand coordination, and an increased speed of wrist movements. This range of motor skills has been investigated using the game Counter Strike, due to its interactive nature. In Counter Strike, players perceive battlefield-like conditions from the first-person perspective, which forces them to engage in various military activities requiring immediate response. The biggest advantage of the top video games players over other opponents is speed, which develops toward expertise.

It should also be added that all subjects in our first study were right-handed, but their left hand, for many years, was extensively used during gaming. It can be seen as intensive training of the left hand, and what we see on the level of VBM are differences in the right hemisphere. Evidence for lateralization related to specific movement has already been well described in the literature (*65, 66*). What is more, the observed result is consistent with the findings of other researchers, who showed that action video game players in comparison to non-players are characterized by faster reaction times in tasks that measure visual and spatial abilities, but only when responses were given using the non-dominant hand (*67*). It should be mentioned that we did not observe a relationship between the size of the lenticular nucleus and experience with RTS games. Other studies (*68*-*70*) have similarly failed to show such correlations, suggesting that the relationship between anatomical plasticity measured using VBM methods and behavior may be more complex and may be mediated by other variables (*71*). There was also low variability in SC2 experience among our participants, which may explain the lack of correlation. The lack of correlation can also be interpreted as an argument for the existence of certain predispositions in complex video game skills acquisition, which we tested and confirmed in our second, training study. Because of the correlational nature of the first study, we cannot determine whether the structural differences between the RTS and NVGP groups were the result of extensive video-game experience or because RTS players have brain structure characteristics that predispose them to engage in activities like playing RTS video games. We designed our longitudinal study, which followed a cross validation procedure (*44*) and introduced a training regimen with the same RTS game as in the first study on a group of NVGPs, to shed some light on this conundrum.

Based on the dominant theory concerning basal ganglia involvement in motor skills we assumed that the differences seen in our first study (cross-sectional) were driven mainly by the motor component of the prolonged SC2 usage. To test that, we followed the methodological approach proposed by Thompson and others (*42*) and focused on the game telemetry. We performed an analysis of both more cognitive game indicators, namely PAC latency, as well as more motor-related game characteristics: APM and HS usage. We found a negative correlation between the GMV of the left putamen and both the left and right pallidum, and PAC latency at the beginning of RTS game skill acquisition. From the cognitive perspective, PACs represent shifts in attention focus followed by a set of motor actions, as SC2 players have to constantly relocate a narrow Point-of-View (PoV) window over a large map area to attend to different locations and execute actions associated with the current state of the game. Technically, each PAC is a PoV that contains one or more actions. PACs encompass roughly 87% of player game time (*42*) and closely resemble the structure of individual trials in experimental tasks that record the set of a participant’s reactions to presented stimuli. We found that PAC latency was relevant to game performance at the beginning of RTS skill acquisition among novice players. This advantage in the early stage of training can be explained by better attentional filtering of relevant game objects. This edge diminishes in later stages of learning, as the game has a finite number of visual elements with meaningful affordances that can be learned over time. For new players, most of the “work” being done is within an individual PAC. To engage a PAC, players have to first attend to a cue, assess what they are looking at within that region of interest, and then start producing actions. PACs latency represents the time it takes for perceptual abilities to paint a picture of the situation, and for attentional abilities to pick through the relevant stimuli. As players accumulate experience and game knowledge, the attentional demands within a PAC should decrease. A larger pretraining basal ganglia volume could boost attention by focally releasing inhibition of task-relevant representations (*72*) at the beginning of learning how to play RTS games. That demand is constantly being stressed through each cycle, and should improve up to some biological limits if game knowledge permits. However, in contrast to our predictions, correlations between HS usage, APM and basal ganglia volume were not found. The usage of HS speeds execution, and the speed of execution is represented by APMs. However, speed plays a very crucial role at the top level of players, but not in novice players and 30 hours of training was likely not enough to develop automatization.

From the perspective of a novice player, success in most RTS video games is by design based on tactical planning, which involves the memory functions and attention of players in many ways. As in almost all strategy games, players devise the most optimal game opening strategies and counter strategies and commit them to memory. For players learning a game like SC2, the most challenging aspect involves memorizing the visuals of interactive game elements and the complex mechanics associated with particular units (e.g. What can that building produce? What types of special actions can this unit make?). Moreover, as the underlying concept of SC2 gameplay is the “counter play” mechanic, it forces players to memorize complex interactions between units (e.g. Which unit will be most effective against a specific threat?), and as the game progresses players need to constantly monitor and update their internal representation of the opponent’s unit composition to react accordingly. On a higher strategic level, RTS video games require players to be able to memorize many game states from their past experience, as this allows for more accurate prediction of the opponents intentions. The putamen and pallidum were shown as uniquely sensitive brain structures in the above-mentioned situations (*73*).

It needs to be added that the obtained results of a greater GMV in the RTS group (our first study) may be interpreted as the effect of a long-term training in the planning and execution of motor sequences (as it has been discussed in the above section), but the GMV of the lenticular nucleus as a neural predictor of RTS training outcomes should be also considered. And the interaction of these two factors seems to be the most probable, as people who engage in intensive and effective video game playing probably have some structural brain predispositions to take such actions and be good at them (which - in turn - motivates them to engage even more). This does not mean that there is no effect of training, but that the correlational nature of our study does not allow us to conclude whether people who start playing video games have different brain structure characteristics in comparison to non-gamers. This unresolved question about brain predispositions in acquiring new skills motivated us to perform a training study.

Our results are in line with Erickson and others (*39*), who showed that putamen volumes were positively correlated with learning new procedures and developing new strategies in a non-commercial RTS video game designed by psychologists. Additionally, Vo and collaborators (*41*) found that patterns of time-averaged T2*- weighted signal in the dorsal striatum recorded before the start of extensive training were predictive of future learning success in the same game. Other regions were recognized as predictors of RTS skill acquisition in an elderly population, such as the prefrontal and frontal regions including the frontal gyrus, anterior commissure, central gyrus, cerebellum, precentral gyrus, and premotor cortex (*40*).

Additionally, we found no effects of training on brain structures in longitudinal study. This result does not support the hypothesis that short term RTS video game training (30h in total) causes alterations in GMV. This does not rule out the possibility that there were changes in GMV, but they are too small to detect using the VBM method (*74*). Using diffusion-weighted MRI to study white matter can provide complementary information about neuroplastic changes after video game training.

## Conclusions and future directions

This paper presents novel findings showing that RTS video game players have a larger lenticular nucleus than NVGPs. Greater volume of the lenticular nucleus can be explained as a result of intensive and complex motor sequence learning (especially automated movements) by our group of RTS experts. However, the contra argument is supported by the assumption that people with specific brain structure characteristics (larger lenticular nucleus) have predispositions to become good video game players. To resolve it, we conducted a second study and checked if there are some neural predispositions that define the manner in which playing a game is learned. We showed that regional differences in volumes of brain areas identified in the first study (on expert RTS players) correlated with the learning pace observed in the second study conducted on a completely new, unrelated and naive to RTS game participants. The present study provides new insights into how skill learning success can depend on brain characteristics. Our results show the importance of individual features of the brain in the effectiveness of training, and in the case of our study - learning to play a new video game. The conclusion that comes to mind is that people with a specific brain structure have a better chance of acquiring new skills. In our study, we showed this in relation to learning a video game, but there is a good chance that it is a more general attribute of the human brain. These findings also point to the usefulness of MRI-brain structure characteristics in predicting relevant intervention outcomes and greatly improve the practicability and effectiveness of those interventions. On the level of a more direct application, our results may open the window to identifying the structural characteristics of successful professional eSports players, much like physical measurements are used in professional sports.

## Materials and Methods - cross-sectional study

### Participants

Sixty-four (n = 64) right-handed, male subjects with a mean age of years 24.55 (SD = 3.66) participated in this study. Two subjects (n = 2) were excluded from the analysis because of bad quality MRI data (image artifacts), so the final sample consisted of sixty-two (n = 62) participants. All subjects completed an on-line questionnaire about demographics, education status, and video-game playing experience. In our self-designed questionnaire (on-line questionnaire on GEX platform) (*75*), we asked additional questions to assess how often individuals engage in various game genres. We broke the games genres down into the following categories: first person shooter (FPS), RTS, role playing, sports, multiplayer online battle arena, racing, puzzles, fighting, turn-based strategy, adventure, and platform games. Inclusion criteria for the RTS experts in our study were as follows: [1] experience with SC2 play, [2] played RTS games at least 6 h/week for the previous six months, [3] declared playing SC2 for more than 60% of total game play time, [4] be an active player (played matches in the last two seasons) and be placed in one of five SC2 leagues (Gold, Platinum, Diamond, Master, Grandmaster). Inclusion criteria for NVGPs were as follows: [1] little or no previous experience with RTS video-game play, and experience with other types of video games totaling no more than 8 h/week (most played less than 6 h) over the past six months. The mean age of the RTS expert group (n = 31) was 24.71 years (SD = 4.27) and 24.39 years (SD = 3.00) in the NVGP group (n = 31).

Education level was matched between groups (all participants were at an undergraduate level). The mean years of education of the RTS expert group was 15.55 years (SD = 2.77) and 16.10 years for the NVGP group (SD = 2.95). We controlled for working memory capacity using the Operation Span Task (OSPAN) (*76*); the mean score was 51.77 (SD = 12.74) for RTS experts and 51.71 (SD = 13.19) for NVGPs. The average hours per week of video games played in different genres from the last six months in RTS experts was 22.74 (11.79) and 2.39 (2.28) for NVGPs. The data from these participants are also a part of our other study (*77*), but the GMV analyses are unpublished in the case of all of the included participants. Table 2 shows the overall video game playing characteristics and average weekly playtime in each video game genre. None of the participants had a history of neurological illness, and they did not declare the use of any psychoactive substances. We also had access to information about each players’ overall performance in the game (wins and losses from the last two seasons) and number of games played.

All subjects participated in additional MRI and cognitive measurement sessions in order to obtain DTI measurements and assess several cognitive functions, which were not related to the project described in this article.

All subjects gave their informed consent to participate in the study, in accordance with the SWPS University Ethical Committee. All participants were male because of difficulties in recruiting female participants with sufficient video game experience. They were paid (approx. 52 USD) for participating in the study.

### MRI - image acquisition

High-resolution whole brain images were acquired on a 3-Tesla MRI scanner (Siemens Magnetom Trio TIM, Erlangen, German) equipped with a 32-channel phased array head coil. T1w images were acquired with the following specification: repetition time, TR = 2530 ms, echo time, TE = 3.32 ms, flip angle, FA = 7°, field of view; FOV = 256 mm, inversion time; TI = 1100 m; voxel size = 1x 1x 1 mm^3^., 176 axial slices. Foam padding was used around the head to minimize head motion during scanning. During these sequences, subjects were asked to relax and try not to fall asleep or move.

The study was a part of a larger project where participants underwent three more MRI sessions (two functional magnetic resonance imaging (fMRI) tasks, diffusion tensor imaging (DTI) session) and a cognitive session on other days.

### Data preprocessing

The same approach was used for both studies. For data preprocessing and statistical analyses, we used Statistical Parametric Mapping (SPM8, Wellcome Trust Center for Neuroimaging, London, UK) running on MATLAB R2015 (The Mathworks, Inc., Natick, MA, USA). We applied standard processing steps as proposed by Ashburner and Friston (2009) (*78*): [1] Checking for anatomical abnormalities and scanner artifacts for each participant, [2] Setting the image origin to the anterior commissure (AC), [3] Manual reorientation to canonical T1 (canonical\avg152T1.nii), [4] Segmentation of tissue classes, [5] Normalization using DARTEL, [6] Modulation of different tissue segments, and [7] Smoothing. A segment algorithm was used in order to obtain basic tissue classes: white matter (WM), gray matter (GM), and cerebrospinal fluid (CSF) (*78*). Next, a study specific template was computed from all participants using Diffeomorphic Anatomical Registration through the Exponentiated Lie Algebra (DARTEL) toolbox (*79*) to determine the nonlinear deformations for warping all the gray and white matter images so that they match each other. This step was followed by affine registration of the gray matter maps to the Montreal Neurological Institute (MNI) space. Modulation (Jacobian determinant) of different tissue segments by nonlinear normalization parameters was applied to correct for individual differences in brain sizes. Finally, data were smoothed with an 8-mm isotropic Gaussian kernel. A group-wise brain mask was computed for statistical analysis to decrease false positives occurring outside the brain. Coordinates of significant effects are reported in MNI space. XjView was used to identify the structures showing effects (http://www.alivelearn.net/xjview). The results were visualized using BrainNet Viewer software (*80*) (http://www.nitrc.org/projects/bnv/).

### Statistical analysis

#### Whole brain GMV comparison between RTS experts and NVGPs

Differences in GMV between RTS experts and NVGPs were calculated using two-sample t-tests. The two-group difference was adjusted for participant age. Given that the total intracranial volume (TIV) could affect the relationships between regional brain volume and measures of skill acquisition, we included TIV in our analyses. An explicit mask was employed (group brain mask with no threshold) to exclude false positives. The group mask was computed by summing GM, WM and CSF for each individual and then computing an average mask for the whole group. The masking was performed using MaskingToolbox (*81*). The model was computed without an absolute threshold since clusters which include voxels with smaller intensity are excluded from the statistical analysis (*81*).

Clusters from the whole brain exploratory analysis (the statistical threshold was set at *p* < 0.001) were corrected to *p* < 0.05 for multiple comparisons using FWE correction at the cluster-level, using a cluster size of 2125 voxels. Next, the average GMV signal from a significant cluster was extracted using the MarsBaR toolbox (*82*). Then, the GMVs (both right and left putamens and pallidums) were fed to the correlation analysis.

## Materials and Methods - training study

### Participants

Twenty subjects (n = 20) participated in the study, but four were excluded from analysis because of low MRI data quality (image artifacts) (n = 2) as well as training dropout (n = 2). The final sample consisted of sixteen (n = 16) right-handed participants with a mean age of 22.94 years (SD = 2.11): five males (22.20 years, SD = 2.39) and eleven females (23.27 years, SD = 2.01). The mean years of education was 15.10 years (SD = 1.93). All subjects completed the same on-line questionnaire as described above (cross-sectional study). Their mean OSPAN score was 52.31 (SD = 18.15). We also asked about their video-game playing experience, and the number of mean weekly hours spent playing video games over the last six months was 0.97 hours (SD = 1.16), with no experience in any action video game genres. None of the participants had a history of neurological illness, and they did not report using psychoactive substances.

All subjects provided written informed consent to participate in the experiment, and the study protocol was approved by the SWPS University Ethical Committee. They were paid (approx. 180 USD) for participating in the study.

### Experimental task

Sixteen participants carried out 30 hours of SC2 gaming in a control laboratory setting. The training lasted from 3 to 4 weeks (a minimum of 6 hours per week, maximum of 10 h per week), with a prohibition of gaming elsewhere (outside the laboratory). Before participants started the training, they had an introduction session with the SC2 trainer. The training was carried out using dedicated desktop PC running Windows 7 (professional edition, 64-bit operating system) equipped with a dedicated graphic card (NVIDIA GeForce GTX 770), 8GB or RAM and a 24” LED display, allowing to play at high graphic quality (1920*1080 pixels resolution, 60Hz). Participants played the game using a mouse/keyboard/headset setup.

In SC2 game players need to build an economy (gathering resources and building bases) and develop the military resources (training units) in order to beat their opponents (destroying their base and army). Cognitive and motor challenges of SC2 game are described in the introduction part of this paper. The participants played using only one Race (Terrans) against AI (artificial intelligence).

There were eight possible difficulty levels in the SC2 matches: Very Easy, Easy, Medium, Hard, Harder, Very Hard, Elite and Cheater. For each victory the player received 1 point. If the player lost, they lost 1 point (−1). The scoring intervals for each level of difficulty were determined as follows: Very Easy - from 0 to 4 points, Easy - from 5 to 8 points, Medium - from 9 to 12 points, Hard - from 13 to 16 points, Harder - from 16 to 20 points, Very Hard - from 21 to 24 points, Elite - from 25 to 28 points, and Cheater - up from 29 points. None of the participants reached the Cheater level, so we included seven levels in the analysis.

We computed the variable indexing the weighted time spent on every level of SC2 difficulty (the time spent on the second level was multiplied by two, the time spent on the third level by three, and so on) for each participant. The final result is a standardized (group-wise) sum of the time spent on all difficulty levels, which reflects performance in the game.

RTS game skill acquisition indicator = (hrs*1)+(hrs*2)+(hrs*3)+(hrs*4)+(hrs*5)+(hrs*6)+(hrs*7)

### MRI - image acquisition

High-resolution T1w images were collected using the IR-FSPGR sequence performed using a 3T MRI GE Discovery MR750w scanner before the RTS training. The MRI scanner was equipped with an 8-channel phased array head coil. T1-weighted (T1w) images were acquired with the following specification: repetition time, TR = 7 ms, echo time, TE = 3 ms, flip angle, FA = 11 °, field of view; FOV= 256 mm, inversion time; TI = 400 ms; voxel size = 1 x 1 x 1 mm^3^, 200 axial slices. Foam padding was used around the head to minimize head motion during scanning. Subjects were asked to try and relax, but not to fall asleep or move. All participants performed a structural MRI and cognitive assessment consisting of several cognitive tasks at two time points: before (T0) and after 30 hours of video game practice (T1). In the current study, we focused on pretraining (T0) MRI scans.

### Data preprocessing

The same approach was used for both studies. Details of data processing are described in Materials and Methods - training study section.

### Statistical analysis

#### Assessing the ROIs for prediction RTS game skill acquisition

Putamen and pallidum were defined using the AAL-116 (*83*) atlas and based on the results from cross-sectional study. Due to the fact that it was a group of novices (non-video game players), we decided to check both putamen and pallidum (bilaterally), and not only within the result obtained from the cross-sectional study where expert video game players were recruited. We aimed to explore this data more due to different skill levels between our groups in the cross-sectional and training study. Each ROI was extracted using the MarsBaR Toolbox (*82*). Then, the GMV was fed to the correlation analysis. Because our data did not meet the assumptions for regression models, all correlational analyses were conducted using Spearman’s correlation coefficient. Correction for multiple comparisons (FDR) for correlation analysis was applied.

## Acknowledgments

**General**: We are grateful to all participants who agreed to be involved in this study. We would like to thank Weronika Debowska for her support during data collection and Michal Chylinski for his help with SC2 participant’s trainings. In Fig. 1. icons were made by Freepik, DinosoftLabs from www.flaticon.com. **Funding:** This study was supported by the Polish National Science Centre NCN grants 2016/23/B/HS6/03843, 2013/10/E/HS6/00186, 2013/11/N/HS6/01335. NK was supported by the Foundation of Polish Science (FNP) and Kosciuszko Foundation.

## Author contributions

NK: designed the study and methods, collected the structural imaging data, collected the behavioral data, analyzed and interpreted data, wrote the manuscript, obtained funding

MS: designed the experiment, SC2 training performed PD: designed the experiment, corrected the manuscript

BK: prepared the MRI sequence and contributed to the interpretation MM: involved in structural imaging data collection

NH: involved in structural imaging data collection MG: involved in behavioral data analysis

AM: involved in structural imaging data analysis, involved in manuscript revising MK: contributed to the interpretation

AB: designed the study and methods, interpreted data, wrote the manuscript, obtained funding

## Data and materials availability

All data needed to evaluate the conclusions in the paper are presented in the paper. Additional data related to this paper and custom analysis scripts are available upon reasonable request.

## References

1. K. Janacsek, J. Fiser, D. Nemeth, The best time to acquire new skills: Age-related differences in implicit sequence learning across the human lifespan. Dev. Sci. 15, 496–505 (2012).

2. D. J. Woltz, An investigation of the role of working memory in procedural skill acquisition. J. Exp. Psychol. Gen. 117, 319 (1988).

3. P. L. Ackerman, R. Kanfer, M. Goff, Cognitive and noncognitive determinants and consequences of complex skill acquisition. J. Exp. Psychol. Appl. 1, 270–304 (1995).

4. O. Matysiak, A. Kroemeke, A. Brzezicka, Working Memory Capacity as a Predictor of Cognitive Training Efficacy in the Elderly Population. Frontiers in Aging Neuroscience. 11 (2019), doi:10.3389/fnagi.2019.00126.

5. C. N. Bürki, C. Ludwig, C. Chicherio, A. de Ribaupierre, Individual differences in cognitive plasticity: an investigation of training curves in younger and older adults. Psychol. Res. 78, 821–835 (2014).

6. J. A. Yesavage, J. I. Sheikh, L. Friedman, E. Tanke, Learning mnemonics: roles of aging and subtle cognitive impairment. Psychol. Aging. 5, 133–137 (1990).

7. M. F. Folstein, S. E. Folstein, P. R. McHugh, “Mini-mental state”: a practical method for grading the cognitive state of patients for the clinician. J. Psychiatr. Res. 12, 189–198 (1975).

8. P. L. Ackerman, A. T. Cianciolo, Cognitive, perceptual-speed, and psychomotor determinants of individual differences during skill acquisition. J. Exp. Psychol. Appl. 6, 259–290 (2000).

9. P. L. Ackerman, Determinants of individual differences during skill acquisition: Cognitive abilities and information processing. J. Exp. Psychol. Gen. 117, 288–318 (1988).

10. J. J. McHenry, L. M. Hough, J. L. Toquam, M. A. Hanson, S. Ashworth, Project A validity results: The relationship between predictor and criterion domains. Pers. Psychol. 43, 335–354 (1990).

11. S. Hutchinson, L. H.-L. Lee, N. Gaab, G. Schlaug, Cerebellar volume of musicians. Cereb. Cortex. 13, 943–949 (2003).

12. S. Tanaka, H. Ikeda, K. Kasahara, R. Kato, H. Tsubomi, S. K. Sugawara, M. Mori, T. Hanakawa, N. Sadato, M. Honda, K. Watanabe, Larger right posterior parietal volume in action video game experts: a behavioral and voxel-based morphometry (VBM) study. PLoS One. 8, e66998 (2013).

13. D. Gong, H. He, D. Liu, W. Ma, L. Dong, C. Luo, D. Yao, Enhanced functional connectivity and increased gray matter volume of insula related to action video game playing. Sci. Rep. 5, 9763 (2015).

14. J. Hänggi, N. Langer, K. Lutz, K. Birrer, S. Mérillat, L. Jäncke, Structural brain correlates associated with professional handball playing. PLoS One. 10, e0124222 (2015).

15. A. Cerasa, A. Sarica, I. Martino, C. Fabbricatore, F. Tomaiuolo, F. Rocca, M. Caracciolo, A. Quattrone, Increased cerebellar gray matter volume in head chefs. PLOS ONE. 12 (2017), p. e0171457.

16. C. Chang, Y. Chen, N. Yen, Nonlinear neuroplasticity corresponding to sports experience: A voxel-based morphometry and resting-state functional connectivity study. Hum. Brain Mapp. 39, 4393–4403 (2018).

17. C. Arkin, E. Przysinda, C. W. Pfeifer, T. Zeng, P. Loui, Gray Matter Correlates of Creativity in Musical Improvisation. Front. Hum. Neurosci. 13, 169 (2019).

18. G. Campitelli, F. Gobet, Deliberate Practice: Necessary But Not Sufficient. Curr. Dir. Psychol. Sci. 20, 280–285 (2011).

19. B. N. Macnamara, D. Z. Hambrick, F. L. Oswald, Deliberate Practice and Performance in Music, Games, Sports, Education, and Professions: A Meta-Analysis. Psychological Science. 25 (2014), pp. 1608–1618.

20. D. Z. Hambrick, A. P. Burgoyne, B. N. Macnamara, F. Ullén, Toward a multifactorial model of expertise: beyond born versus made. Annals of the New York Academy of Sciences. 1423 (2018), pp. 284–295.

21. E. A. Maguire, Navigation-related Structural Change in the Hippocampi of Taxi Drivers (2000).

22. N. Golestani, C. J. Price, S. K. Scott, Born with an ear for dialects? Structural plasticity in the expert phonetician brain. J. Neurosci. 31, 4213–4220 (2011).

23. C. Thomas, C. I. Baker, Teaching an adult brain new tricks: a critical review of evidence for training-dependent structural plasticity in humans. Neuroimage. 73, 225– 236 (2013).

24. B. Draganski, C. Gaser, V. Busch, G. Schuierer, U. Bogdahn, A. May, Neuroplasticity: changes in grey matter induced by training. Nature. 427, 311–312 (2004).

25. S. Kühn, T. Gleich, R. C. Lorenz, U. Lindenberger, J. Gallinat, Playing Super Mario induces structural brain plasticity: gray matter changes resulting from training with a commercial video game. Mol. Psychiatry. 19, 265–271 (2014).

26. M. Palaus, E. M. Marron, R. Viejo-Sobera, D. Redolar-Ripoll, Neural Basis of Video Gaming: A Systematic Review. Front. Hum. Neurosci. 11, 248 (2017).

27. J. Legault, A. Grant, S.-Y. Fang, P. Li, A longitudinal investigation of structural brain changes during second language learning. Brain Lang. 197, 104661 (2019).

28. K. C. Barrett, R. Ashley, D. L. Strait, N. Kraus, Art and science: how musical training shapes the brain. Front. Psychol. 4, 713 (2013).

29. C. E. James, M. S. Oechslin, D. Van De Ville, C.-A. Hauert, C. Descloux, F. Lazeyras, Musical training intensity yields opposite effects on grey matter density in cognitive versus sensorimotor networks. Brain Struct. Funct. 219, 353–366 (2014).

30. K. I. Erickson, M. W. Voss, R. S. Prakash, C. Basak, A. Szabo, L. Chaddock, J. S. Kim, S. Heo, H. Alves, S. M. White, T. R. Wojcicki, E. Mailey, V. J. Vieira, S. A. Martin, B. D. Pence, J. A. Woods, E. McAuley, A. F. Kramer, Exercise training increases size of hippocampus and improves memory. Proc. Natl. Acad. Sci. U. S. A. 108, 3017–3022 (2011).

31. L. Cui, H. Yin, S. Lyu, Q. Shen, Y. Wang, X. Li, J. Li, Y. Li, L. Zhu, Tai Chi Chuan vs General Aerobic Exercise in Brain Plasticity: A Multimodal MRI Study. Sci. Rep. 9, 17264 (2019).

32. M. Lövdén, L. Bäckman, U. Lindenberger, S. Schaefer, F. Schmiedek, A theoretical framework for the study of adult cognitive plasticity. Psychol. Bull. 136, 659–676 (2010).

33. E. Wenger, C. Brozzoli, U. Lindenberger, M. Lövdén, Expansion and Renormalization of Human Brain Structure During Skill Acquisition. Trends Cogn. Sci. 21, 930–939 (2017).

34. N. Lehmann, J. W. Tolentino-Castro, E. Kaminski, P. Ragert, A. Villringer, M. Taubert, Interindividual differences in gray and white matter properties are associated with early complex motor skill acquisition. Hum. Brain Mapp. 40, 4316–4330 (2019).

35. K. R. Sherrill, E. R. Chrastil, I. Aselcioglu, M. E. Hasselmo, C. E. Stern, Structural Differences in Hippocampal and Entorhinal Gray Matter Volume Support Individual Differences in First Person Navigational Ability. Neuroscience. 380, 123– 131 (2018).

36. S. Park, S.-H. Ryu, Y. Yoo, J.-J. Yang, H. Kwon, J.-H. Youn, J.-M. Lee, S.-J. Cho, J.-Y. Lee, Neural predictors of cognitive improvement by multi-strategic memory training based on metamemory in older adults with subjective memory complaints. Sci. Rep. 8, 1095 (2018).

37. E. Moore, R. Schaefer, M. Bastin, N. Roberts, K. Overy, Can Musical Training Influence Brain Connectivity? Evidence from Diffusion Tensor MRI. Brain Sciences. 4 (2014), pp. 405–427.

38. D. Momi, C. Smeralda, G. Sprugnoli, S. Ferrone, S. Rossi, A. Rossi, G. Di Lorenzo, E. Santarnecchi, Acute and long-lasting cortical thickness changes following intensive first-person action videogame practice. Behav. Brain Res. 353, 62–73 (2018).

39. K. I. Erickson, W. R. Boot, C. Basak, M. B. Neider, R. S. Prakash, M. W. Voss, A. M. Graybiel, D. J. Simons, M. Fabiani, G. Gratton, A. F. Kramer, Striatal volume predicts level of video game skill acquisition. Cereb. Cortex. 20, 2522–2530 (2010).

40. C. Basak, M. W. Voss, K. I. Erickson, W. R. Boot, A. F. Kramer, Regional differences in brain volume predict the acquisition of skill in a complex real-time strategy videogame. Brain Cogn. 76, 407–414 (2011).

41. L. T. K. Vo, D. B. Walther, A. F. Kramer, K. I. Erickson, W. R. Boot, M. W. Voss, R. S. Prakash, H. Lee, M. Fabiani, G. Gratton, D. J. Simons, B. P. Sutton, M. Y. Wang, Predicting individuals’ learning success from patterns of pre-learning MRI activity. PLoS One. 6, e16093 (2011).

42. J. J. Thompson, M. R. Blair, L. Chen, A. J. Henrey, Video game telemetry as a critical tool in the study of complex skill learning. PLoS One. 8, e75129 (2013).

43. D. Bavelier, B. Bediou, C. Shawn Green, Expertise and generalization: lessons from action video games. Current Opinion in Behavioral Sciences. 20 (2018), pp. 169– 173.

44. F. Hoeft, T. Ueno, A. L. Reiss, A. Meyler, S. Whitfield-Gabrieli, G. H. Glover, T. A. Keller, N. Kobayashi, P. Mazaika, B. Jo, M. A. Just, J. D. E. Gabrieli, Prediction of children’s reading skills using behavioral, functional, and structural neuroimaging measures. Behav. Neurosci. 121, 602–613 (2007).

45. F. Faul, E. Erdfelder, A. Buchner, A.-G. Lang, Statistical power analyses using G*Power 3.1: tests for correlation and regression analyses. Behav. Res. Methods. 41, 1149–1160 (2009).

46. A. M. Graybiel, Basal ganglia—input, neural activity, and relation to the cortex. Curr. Opin. Neurobiol. 1, 644–651 (1991).

47. J. L. Lanciego, N. Luquin, J. A. Obeso, Functional Neuroanatomy of the Basal Ganglia. Cold Spring Harbor Perspectives in Medicine. 2 (2012), pp. a009621– a009621.

48. H. Boecker, A. Dagher, A. O. Ceballos-Baumann, R. E. Passingham, M. Samuel, K. J. Friston, J. Poline, C. Dettmers, B. Conrad, D. J. Brooks, Role of the human rostral supplementary motor area and the basal ganglia in motor sequence control: investigations with H2 15O PET. J. Neurophysiol. 79, 1070–1080 (1998).

49. R. S. Turner, S. T. Grafton, J. R. Votaw, M. R. Delong, J. M. Hoffman, Motor Subcircuits Mediating the Control of Movement Velocity: A PET Study. Journal of Neurophysiology. 80 (1998), pp. 2162–2176.

50. W. R. Marchand, J. N. Lee, J. W. Thatcher, E. W. Hsu, E. Rashkin, Y. Suchy, G. Chelune, J. Starr, S. S. Barbera, Putamen coactivation during motor task execution. Neuroreport. 19, 957–960 (2008).

51. R. S. Turner, M. Desmurget, J. Grethe, M. D. Crutcher, S. T. Grafton, Motor subcircuits mediating the control of movement extent and speed. J. Neurophysiol. 90, 3958–3966 (2003).

52. M. Corbetta, F. M. Miezin, S. Dobmeyer, G. L. Shulman, S. E. Petersen, Selective and divided attention during visual discriminations of shape, color, and speed: functional anatomy by positron emission tomography. J. Neurosci. 11, 2383– 2402 (1991).

53. F. McNab, T. Klingberg, Prefrontal cortex and basal ganglia control access to working memory. Nat. Neurosci. 11, 103–107 (2008).

54. Y. V. Martos, B. Y. Braz, J. P. Beccaria, M. G. Murer, J. E. Belforte, Compulsive Social Behavior Emerges after Selective Ablation of Striatal Cholinergic Interneurons. J. Neurosci. 37, 2849–2858 (2017).

55. M. van Beilen, K. L. Leenders, Putamen FDOPA uptake and its relationship tot cognitive functioning in PD. J. Neurol. Sci. 248, 68–71 (2006).

56. N. Arimura, Y. Nakayama, T. Yamagata, J. Tanji, E. Hoshi, Involvement of the Globus Pallidus in Behavioral Goal Determination and Action Specification. Journal of Neuroscience. 33 (2013), pp. 13639–13653.

57. J. J. Thompson, C. M. McColeman, E. R. Stepanova, M. R. Blair, Using video game telemetry data to research motor chunking, action latencies, and complex cognitive-motor skill learning. Top. Cogn. Sci. 9, 467–484 (2017).

58. J. J. Thompson, C. M. McColeman, M. R. Blair, A. J. Henrey, Classic motor chunking theory fails to account for behavioural diversity and speed in a complex naturalistic task. PLoS One. 14, e0218251 (2019).

59. D. Bavelier, B. Bediou, C. S. Green, Expertise and generalization: lessons from action video games. Current Opinion in Behavioral Sciences. 20, 169–173 (2018).

60. D. Bavelier, B. Bediou, C. Shawn Green, Expertise and generalization: lessons from action video games. Current Opinion in Behavioral Sciences. 20 (2018), pp. 169– 173.

61. I. H. Jenkins, D. J. Brooks, P. D. Nixon, R. S. Frackowiak, R. E. Passingham, Motor sequence learning: a study with positron emission tomography. The Journal of Neuroscience. 14 (1994), pp. 3775–3790.

62. S. Lehericy, H. Benali, P.-F. Van de Moortele, M. Pelegrini-Issac, T. Waechter, K. Ugurbil, J. Doyon, Distinct basal ganglia territories are engaged in early and advanced motor sequence learning. Proceedings of the National Academy of Sciences. 102 (2005), pp. 12566–12571.

63. D. Coynel, G. Marrelec, V. Perlbarg, M. Pélégrini-Issac, P.-F. Van de Moortele, K. Ugurbil, J. Doyon, H. Benali, S. Lehéricy, Dynamics of motor-related functional integration during motor sequence learning. Neuroimage. 49, 759–766 (2010).

64. L. Borecki, K. Tolstych, M. Pokorski, Computer games and fine motor skills. Adv. Exp. Med. Biol. 755, 343–348 (2013).

65. R. Kawashima, K. Yamada, S. Kinomura, T. Yamaguchi, H. Matsui, S. Yoshioka, H. Fukuda, Regional cerebral blood flow changes of cortical motor areas and prefrontal areas in humans related to ipsilateral and contralateral hand movement. Brain Res. 623, 33–40 (1993).

66. A. Solodkin, P. Hlustik, D. C. Noll, S. L. Small, Lateralization of motor circuits and handedness during finger movements. European Journal of Neurology. 8 (2001), pp. 425–434.

67. F. Richlan, J. Schubert, R. Mayer, F. Hutzler, M. Kronbichler, Action video gaming and the brain: fMRI effects without behavioral effects in visual and verbal cognitive tasks. Brain Behav. 8, e00877 (2018).

68. S. Tanaka, H. Ikeda, K. Kasahara, R. Kato, H. Tsubomi, S. K. Sugawara, M. Mori, T. Hanakawa, N. Sadato, M. Honda, K. Watanabe, Larger right posterior parietal volume in action video game experts: a behavioral and voxel-based morphometry (VBM) study. PLoS One. 8, e66998 (2013).

69. J. Boyke, J. Driemeyer, C. Gaser, C. Büchel, A. May, Training-induced brain structure changes in the elderly. J. Neurosci. 28, 7031–7035 (2008).

70. Ł. Bola, M. Zimmermann, P. Mostowski, K. Jednoróg, A. Marchewka, P. Rutkowski, M. Szwed, Task-specific reorganization of the auditory cortex in deaf humans. Proc. Natl. Acad. Sci. U. S. A. 114, E600–E609 (2017).

71. C. Sampaio-Baptista, J. Scholz, M. Jenkinson, A. G. Thomas, N. Filippini, G. Smit, G. Douaud, H. Johansen-Berg, Gray matter volume is associated with rate of subsequent skill learning after a long term training intervention. Neuroimage. 96, 158– 166 (2014).

72. M. R. van Schouwenburg, H. E. M. den Ouden, R. Cools, Selective attentional enhancement and inhibition of fronto-posterior connectivity by the basal ganglia during attention switching. Cereb. Cortex. 25, 1527–1534 (2015).

73. S. W. Ell, S. Helie, S. Hutchinson, A. Costa, E. Villalba, Contributions of the putamen to cognitive function. Horizons in neuroscience research (2011), pp. 29–52.

74. R. J. Zatorre, R. D. Fields, H. Johansen-Berg, Plasticity in gray and white: neuroimaging changes in brain structure during learning. Nat. Neurosci. 15, 528–536 (2012).

75. B. Sobczyk, P. Dobrowolski, M. Skorko, J. Michalak, A. Brzezicka, Issues and advances in research methods on video games and cognitive abilities. Front. Psychol. 6, 1451 (2015).

76. N. Unsworth, R. P. Heitz, J. C. Schrock, R. W. Engle, An automated version of the operation span task. Behav. Res. Methods. 37, 498–505 (2005).

77. N. Kowalczyk, F. Shi, M. Magnuski, M. Skorko, P. Dobrowolski, B. Kossowski, A. Marchewka, M. Bielecki, M. Kossut, A. Brzezicka, Real-time strategy video game experience and structural connectivity - A diffusion tensor imaging study. Hum. Brain Mapp. 39, 3742–3758 (2018).

78. J. Ashburner, K. J. Friston, Voxel Based Morphometry. Encyclopedia of Neuroscience (2009), pp. 471–477.

79. J. Ashburner, A fast diffeomorphic image registration algorithm. Neuroimage. 38, 95–113 (2007).

80. M. Xia, J. Wang, Y. He, BrainNet Viewer: a network visualization tool for human brain connectomics. PLoS One. 8, e68910 (2013).

81. G. Ridgway, R. Omar, S. Ourselin, D. Hill, J. Warren, N. Fox, Issues with threshold masking in voxel-based morphometry of atrophied brains. NeuroImage. 44 (2009), pp. 99–111.

82. M. Brett, J.-L. Anton, R. Valabregue, J.-B. Poline, Others, in 8th international conference on functional mapping of the human brain (Sendai, Japan, 2002), vol. 16, p. 497.

83. N. Tzourio-Mazoyer, B. Landeau, D. Papathanassiou, F. Crivello, O. Etard, N. Delcroix, B. Mazoyer, M. Joliot, Automated anatomical labeling of activations in SPM using a macroscopic anatomical parcellation of the MNI MRI single-subject brain. Neuroimage. 15, 273–289 (2002).

