## Supplemental Table 1 for "Lenticular nucleus volume predicts performance in real-time strategy game - cross-sectional and training approach using voxel-based morphometry"

**Table 1. Correlation coefficient (*r*) from Spearman's correlation analyses between ROIs GMW (left, right putamen and pallidum) and Perception Action Cycle latency, Actions Per Minute, Hotkey Selects usage.**

| | Putamen L | Putamen R | Pallidum L | Pallidum R |
|---|---|---|---|---|
| PAC latency Q1 | ***r* = -0.58, *p* = 0.02** | *r* = -0.43, *p* = 0.10 | ***r* = -0.57, *p* = 0.02** | ***r* = 0.54, *p* = 0.03** |
| PAC latency Q2 | *r* = -0.34, *p* = 0.20 | *r* = -0.12, *p* = 0.66 | *r* = -0.33, *p* = 0.21 | *r* = -0.25, *p* = 0.35 |
| PAC latency Q3 | *r* = -0.20, *p* = 0.46 | *r* = -0.10, *p* = 0.72 | *r* = -0.26, *p* = 0.34 | *r* = -0.22, *p* = 0.41 |
| PAC latency Q4 | *r* = -0.21, *p* = 0.43 | *r* = -0.19, *p* = 0.49 | *r* = -0.30, *p* = 0.26 | r = -0.29, *p* = 0.27 |
| APM Q1 | *r* = 0.22, *p* = 0.41 | *r* = 0.12, *p* = 0.65 | *r* = 0.03, *p* = 0.91 | *r* = -0.04, *p* = 0.88 |
| APM Q2 | *r* = 0.16, *p* = 0.56 | *r* = -0.02, *p* = 0.94 | *r* = -0.03, *p* = 0.91 | *r* = -0.06, *p* = 0.83 |
| APM Q3 | *r* = 0.23, *p* = 0.39 | *r* = 0.09, *p* = 0.75 | *r* = 0.06, *p* = 0.82 | *r* = -0.03, *p* = 0.91 |
| APM Q4 | *r* = 0.25, *p* = 0.36 | *r* = 0.09, *p* = 0.74 | *r* = 0.10, *p* = 0.71 | *r* = 0.06, *p* = 0.82 |
| HS Q1 | *r* = -0.36, *p* = 0.18 | *r* = -0.40, *p* = 0.13 | *r* = -0.29, *p* = 0.28 | *r* = -0.39, *p* = 0.13 |
| HS Q2 | *r* = -0.27, *p* = 0.31 | *r* = -0.24, *p* = 0.38 | *r* = -0.14, *p* = 0.59 | *r* = -0.27, *p* = 0.32 |
| HS Q3 | *r* = -0.28, *p* = 0.28 | *r* = -0.31, *p* = 0.25 | *r* = -0.20, *p* = 0.45 | *r* = -0.33, *p* = 0.21 |
| HS Q4 | *r* = -0.18, *p* = 0.51 | *r* = -0.18, *p* = 0.51 | *r* = -0.07, *p* = 0.79 | *r* = -0.21, *p* = 0.44 |

*Significant results are bolded, uncorrected for multiple comparisons (false discovery rate). PAC - Perception Action Cycle, APM - Action Per Minute, HS - Hotkey Selects, Q - quartile, L - left, R – right
