## Supplemental Table 2 for "Lenticular nucleus volume predicts performance in real-time strategy game - cross-sectional and training approach using voxel-based morphometry"

**Table 2. Mean weekly hours of video games played in the last 6 months, by genre.** Values are mean and SD (SD in parentheses). All variables were compared between groups with two sample t-tests**.**

| Video game genre | RTS experts | NVGPs | *p*-value |
| --- | --- | --- | --- |
| Real-time strategy | **16.06 (9.91)** | **0.05 (0.15)** | **0.000** |
| First-person shooter | 1.02 (2.15) | 0.27 (0.60) | 0.07 |
| Platform | 0 (0.00) | 0.06 (0.21) | 0.10 |
| Fighting | 0.16 (0.57) | 0 (0.00) | 0.12 |
| Turn-based strategy | 1 (1.77) | 0.35 (0.70) | 0.06 |
| Sports | **0 (0.00)** | **0.65 (1.42)** | **0.01** |
| Role-play | **1.60 (2.08)** | **0.19 (0.46)** | **0.01** |
| Racing | 0.13 (0.29) | 0.27 (0.55) | 0.20 |
| Logic | 0.66 (1.32) | 0.37 (0.66) | 0.30 |
| Multiplayer online battle arena | **1.44 (2.99)** | **0.13 (0.39)** | **0.02** |
| Adventure | 0.68 (1.88) | 0.03 (0.12) | 0.06 |

*Significant results are bolded. RTS - real time strategy games, NVGPs – non-video game players

The data presented in table 2 are also a part of our other study (*77*).
